# Inducible expression of Oct-3/4 reveals synergy with Klf4 in targeting Cyclin A2 to enhance proliferation during early reprogramming

**DOI:** 10.1101/2021.03.16.435687

**Authors:** Lamuk Zaveri, Jyotsna Dhawan

**Author notes:** Address correspondence to: **Jyotsna Dhawan**, Centre for Cellular and Molecular Biology, Uppal Road Hyderabad 500 007 India.

## Abstract

During reprogramming of somatic cells, heightened proliferation is one of the earliest changes observed. While other early events such as mesenchymal-to-epithelial transition have been well studied, the mechanisms by which the cell cycle switches from a slow cycling state to a faster cycling state are still incompletely understood. To investigate the role of Oct-3/4 in this early feature of reprogramming, we created a 4-Hydroxytamoxifen dependent Oct-3/4 Estrogen Receptor fusion (OctER). We show that OctER can substitute for Oct-3/4 to reprogram mouse embryonic fibroblasts to induced pluripotent stem cells. While over-expression of OctER or Klf4 individually did not affect cell proliferation, in combination, these factors hasten the cell cycle, in a tamoxifen dose-dependent manner, supporting a key role for OctER. Oct-3/4 + Klf4 increased proliferation by enhancing expression of Cyclin A2. We verified occupancy of endogenous Oct-3/4 and Klf4 at bioinformatically identified binding sites in the Cyclin A2 promoter in mouse embryonic stem cells (mESC). Using inducible OctER along with Klf4, we show dose-dependent induction of Cyclin A2 promoter-reporter activity and mRNA levels. Taken together, our results provide further evidence of the interdependence of pluripotency and the rapid cell cycle seen in mESC, and identify CyclinA2 as a key early target.

## Introduction

During embryogenesis, there is an enormous requirement for new cells derived from early pluripotent stem cells (PSC). Compared to somatic cells, PSC exhibit a modified cell cycle, allowing for extremely rapid proliferation, keeping pace with enlargement and specialisation of the growing embryo [1–3]. The rapid proliferation is enabled by a major rearrangement of cell cycle regulators, such that ESC primarily oscillate between DNA synthesis (S phase) and mitosis (M phase) spaced by a highly reduced G1 phase [1, 2]. ESC undergo an estimated 40 divisions during embryonic mouse development, while somatic adult progenitors may divide a total of 5 times during the subsequent lifetime of the animal, and most differentiated cells never divide following specialization. Thus, rapid proliferation is tightly linked to the pluripotent state.

Mouse ESC (mESC) derived from the inner cell mass of the blastocyst have a doubling time of 8 to 10 hours in vitro and 4.5 to 7.5 hours in vivo [4–6]. A prominent feature of this rapid cell cycle is reduced activity of retinoblastoma protein (RB), gatekeeper of the G1-S phase transition [7]. High activity and non-oscillatory expression of Cyclin A2 and Cyclin E, and their partner Cdk2, keep RB in an inactive hyperphosphorylated state [1, 2]. Absence of cell cycle inhibitors such as the cyclin dependent kinase inhibitor (CDKI) p21 further ensures that mESC transit G1-S more rapidly [1, 2]. PSC progression to multi-potent and finally mono-potent cells is accompanied by lengthening of the cell cycle, resulting in reduced proliferation rates [3]. Cell cycle extension is mainly due to increased G1 duration, by activation of RB, and the CDKIs [3, 5]. The molecular mechanisms that regulate cell cycle duration during development are not completely understood.

ESCs are a useful system to dissect these mechanisms, due to their ability to differentiate to most cell types, coupled with their limitless ability to self-renew. Given the extensive links between pluripotency and rapid cell cycle, uncovering the role of pluripotency factors in cell cycle control is more readily addressed during reprogramming of somatic cells [8]. The slower proliferation and longer G1 of somatic cells switches to the faster cell cycle seen in mESC upon forced expression of the Yamanaka factors [9]. Cell cycle shortening is initiated early, when exogenous Yamanaka factors dominate, permitting an analysis of how these factors affect the cell cycle during the gain of pluripotency [9–12]. Of the 4 Yamanaka factors, c-Myc is dispensable for reprogramming and Sox2 is required for the later stages of reprogramming [13–16], suggesting that Klf4 and Oct-3/4 are likely to regulate early cell cycle changes during reprogramming.

Of the two transcription factors, the role of Klf4 is complicated as it can function either as transcriptional activator or repressor, with functional redundancy amongst Klf family members [17, 18]. However, Oct-3/4 has been shown to play a role in maintaining the rapid ESC cell cycle [5], and Oct-3/4 levels are very tightly maintained [19]. Thus, Oct-3/4 may play a major role in regulating the early cell cycle alterations during reprogramming.

To investigate the role of Oct-3/4 in the early stages of reprogramming, we generated a 4-Hydroxytamoxifen (OHT) inducible Oct-3/4 (OctER). Using OctER, we reprogrammed primary mouse embryonic fibroblasts (MEF) to a pluripotent state in conjunction with Klf4, Sox2 and c-Myc. The induced pluripotent stem cells (iPSC) thus produced were found to exhibit hallmarks of pluripotency. Since OctER could substitute for Oct-3/4 in reprogramming, we used the inducible factor to assess cell cycle changes that occur during the early stages of reprogramming. We show that Oct-3/4 in conjunction with Klf4 enhances the early rise in expression of Cyclin A2, a key positive cell cycle regulator of the G1-S transition, and thereby enhances MEF proliferation towards levels essential for reprogramming.

## Materials and Methods

### Plasmids

Oct-3/4 (from pCX-OKS-2A) was expressed from a CMV promoter in a modified FUGW plasmid; to generate OctER, the above Oct-3/4 construct was engineered with a modified OHT binding domain from pBABE-cMycER. For Klf4, pLOVE-Klf4 was used. Mouse Nanog promoter luciferase reporter was from Dr. Tada. Cyclin A2 promoter (−500bp to +209bp) was cloned from C57BL/6J mouse genomic DNA into pGL3 basic vector. Primers and plasmids used are in Table S3, S4. Addgene plasmids are acknowledged.

### Luciferase assay

Cos7 cells were transfected with Nanog promoter-luc or Cyclin A2 promoter-luc along with pRL-SV40 (Promega, USA), Klf4 or Oct-3/4 or Klf4 + Oct-3/4 or Klf4 + OctER using PEI (Polysciences, USA). Self-ligated empty pGEM-T vector (Promega, USA) was used to ensure equal amounts of transfected DNA. 24 hours post transfection, fresh media was added with OHT or ethanol (vehicle control). 24 hours post induction, cells were lysed with Passive Lysis Buffer (PLB) from the Dual-Luciferase Reporter Assay (Promega, USA), Luciferase Assay Reagent (LARII) was added to measure firefly luciferase activity (F-luc) after which Stop & Glo was added to measure renilla luciferase activity (R-luc) on an EnSpire Multimode Plate Reader (Perkin Elmer, USA). F-luc was normalised to R-luc to account for transfection efficiency, and normalised to the respective basal promoter activity.

### Isolating nuclear fractions

Nuclear fractions were isolated using the REAP method [49]. MEFs expressing OctER were pulsed with either OHT or ethanol, 24 hours later were washed in PBS, collected by scraping into 1.5 ml ice cold PBS + protease inhibitors (Roche, Switzerland), flash spun, resuspended in 180 μl chilled 0.1% NP40. 90 μl of whole cell lysate (WCL) was set aside; the remainder was flash spun and 90 μl of supernatant saved as cytoplasmic fraction. The pellet was resuspended in 90 μl chilled 0.1% NP40, flash spun, supernatant discarded and the pellet (nuclear fraction) was resuspended in 1X Laemmli buffer. 4X Laemmli buffer was added to WCL and cytoplasmic fractions. All samples were briefly sonicated (Bioruptor, Diagenode Belgium) and boiled before loading on SDS-PAGE.

### Western blotting

Equal volumes of nuclear fraction (equal cell numbers) or 20 μg of protein was resolved by 8 % SDS-PAGE, transferred to PVDF membrane (Millipore, USA), blocked with Blotto (Santa Cruz, USA) and probed with antibodies against Oct-3/4 (Santa Cruz, USA), Gapdh (Abcam, USA) or Lamin B1 (Abcam, USA). The membrane was washed and probed with HRP-conjugated secondary antibodies (Jackson, USA), proteins visualised using ECL detection reagent (Amersham, USA) and imaged using ChemiDoc (Syngene, USA). Densitometric analysis was performed using ImageJ. The intensity of OctER was normalised to loading control Lamin B1 and plotted in arbitrary units with respect to uninduced.

### RNA isolation and qRT-PCR

RNA from MEFs were isolated at different time points post-transduction using RNeasy Micro kit (Qiagen, Germany). RNA from mESC was isolated using TRIzol. For iPSC, cells were passaged feeder free for three passages before RNA was isolated using TRIzol. 1-2 μg total RNA was used for first strand synthesis using Oligo (dT) and SuperScript III First-Strand Synthesis System. qRT-PCR was performed on an ABI ViiA 7 or 7900HT using Power SYBR Green PCR Master Mix. Ct values were first normalised to Gapdh to calculate ΔCt. ΔΔCt was calculated with respect to either untransduced MEFs or MEFs transduced with empty vector or uninduced in the case of OHT treatment. Average of 3 independent biological replicates was plotted as 2^-ΔΔCt^ to calculate fold change with standard error using error propagation. Primer sequences used for qRT-PCR are provided in Table S5. All reagents (except RNeasy) and instruments were from ThermoScientific, USA.

### MEF derivation

All mouse experiments were approved by the inStem/NCBS or CCMB Institutional Animal Ethics Committee and performed under CPCSEA guidelines.

A pregnant female Oct-eGFP (OG-2) mouse at 13.5 dpc was sacrificed by cervical dislocation and the embryos were harvested, washed well in PBS, the head and liver removed and remaining tissue collected in PBS, minced and homogenized by syringing. Tissue slurry was enzymatically digested, and cells harvested by centrifugation and plated for expansion till 95% confluency, when they were trypsinised and cryopreserved as P0. MEFs were used till P4.

### Cell culture

MEFs, Cos7, Lenti-X 293T were maintained in DMEM high glucose, 10% heat inactivated FBS, 10,000 units/ml penicillin, 10 mg/ml streptomycin, 1 mM sodium pyruvate, Glutamax. mESC E14 and iPSC were maintained in DMEM/F12, 15% KnockOUT serum replacement, NEAA, LIF, β-Mercaptoethanol, 10,000 units/ml penicillin, 10 mg/ml streptomycin, Glutamax. All culture components were from ThermoScientific, USA or Sigma, USA.

### Virus production and transduction

All viral work was approved by inStem or CCMB Institutional Biosafety Committees. Lenti-X 293T cells (TakaraBio, USA) were transfected with lentiviral expression vector, psPAX2 and pHCMV-EcoEnv in the molar ratio of 2:1:1 using PEI. Culture supernatants containing virus were harvested 24, 48, 72 hours post transfection, pooled, cleared by centrifugation, filtered through a 0.45μm PES filter (Nalgene, USA), concentrated 10 fold using Amicon Ultra-15 centrifugal filter unit, 100MWCO (Millipore, USA) and filtered through a 0.45μm PES filter (Nalgene, USA). MEFs were transduced with concentrated viral supernatant with 8 μg/μl Polybrene (Sigma, USA).

### Reprogramming

All stem cell work was approved by the inStem or CCMB Institutional Committee on Stem Cell Research. MEFs from OG-2 mice were used for reprogramming from P1-4, by transduction with viral supernatant containing equal amounts of either Oct-3/4 + Klf4 + Sox2 + c-Myc or OctER + Klf4 + Sox2 + c-Myc along with Polybrene. 48 hours post transduction, MEFs were plated on gelatin coated plates containing Mitomycin-C inactivated feeders. Cells were pulsed daily with OHT in fresh media; media was switched to ESC cell media on Day 2. To calculate reprogramming efficiency, 5 x 10^5^ transduced MEFs were seeded on feeders in duplicate. Reprogramming was carried out for 21 days, cells were fixed and stained for Alkaline Phosphatase activity using Leukocyte Alkaline Phosphatase Kit (Sigma, USA). Reprogramming efficiency was calculated as a percentage of compact alkaline phosphatase colonies to cells seeded.

### Immunostaining

MEFs expressing OctER were seeded onto coverslips, pulsed with OHT after 12 hours, fixed after 24 hours with 4% PFA, blocked, permeabilised and incubated with antibody against Oct-3/4 (Santa Cruz, USA). Cells were washed and incubated with Alexa-488 conjugated secondary antibody (ThermoScientific, USA) and DAPI (Sigma, USA). The coverslips were mounted using glycerol and imaged on an Axio Imager Z2 microscope (Zeiss, Germany). iPSC were seeded onto gelatin-coated coverslips and fixed after 24 hours, blocked, permeabilised and incubated with antibodies against SSEA-1 (Millipore, USA) or Nanog (Abcam, USA), washed and incubated with Alexa-546 conjugated secondary antibody (ThermoScientific, USA) and DAPI (Sigma, USA). The coverslips were mounted using glycerol and imaged on SP5 confocal microscope (Leica, Germany).

### Teratoma assay and histological analysis

All mouse experiments were approved by the CCMB Institutional Animal Ethics committee and performed under CPCSEA guidelines. A cell suspension of 5 x 10^6^ cells in 200 μl PBS was injected sub-cutaneously into nude mice. 4 weeks later, mice were sacrificed by cervical dislocation, teratoma surgically excised, measured and imaged, and a fragment embedded in paraffin, sectioned using a Leica microtome, stained using Hematoxylin & Eosin and imaged on an Axio Imager Z2 microscope (Zeiss, Germany).

### WST assay

Transduced MEFs were seeded in triplicate in a 96 well flat bottom tissue culture plate and WST-1 assay (Roche, Switzerland) performed as per manufacturer’s protocol at day 0, 1, 3 and 5. Absorbance was measured at 450 nm and 640 nm. Background corrected values were plotted with respect to Day 0.

### Bioinformatic analysis of published data sets

Datasets in BAM, BED, or bigWig format were downloaded from NCBI GEO, processed on Galaxy (https://usegalaxy.org/) and visualised on the UCSC genome browser (http://genome.ucsc.edu/index.html). Enrichment peaks were identified across the loci of cell cycle genes, including 5 kb upstream of TSS.

### ChIP

mESC were trypsinised, counted and fixed in a 1% formaldehyde-PBS for 10 minutes at room temperature. Formaldehyde was quenched with 1.25 M Glycine for 5 minutes at room temperature, cells were washed 3 times with PBS, the cell pellet snap-frozen and stored at −80°C. Cell pellets were thawed on ice in SDS-lysis solution (ChIP assay kit, Millipore, USA) + protease inhibitors for 15 minutes with intermittent vortexing to ensure complete lysis. Chromatin was sonicated using a Bioruptor Pico (Diagenode, Belgium) for 25 cycles with 15 seconds on, 45 seconds off, centrifuged for 10 minutes and supernatant transferred to a fresh tube. ChIP was performed as per manufacturer’s instructions (Millipore, USA). 5 μg of polyclonal anti-Oct-3/4 (Santa Cruz, USA), polyclonal anti-Klf4 (R & D Systems, USA) or Goat IgG (Jackson, USA) was used per 1.5 x 10^6^ cells. DNA was purified using MinElute PCR purification Kit (Qiagen, Germany). qRT-PCR was performed with purified DNA using primers listed in Table S6. Enrichment was calculated as % input.

### Statistical analysis

Data shown is the average of at least three independent experiments along with standard error. To calculate P value, Student’s t test was used. All statistical analyses was done in MS Excel.

## Results

### Constructing an OHT dependent inducible Oct-3/4 (OctER)

To generate stable expression constructs for inducible Oct-3/4, we used a lentiviral system. Oct-3/4 protein consists of two DNA-binding domains linked by a flexible linker that permit binding as a monomer, a homodimer or a heterodimer with other proteins, such as Sox2 [20]. To preserve this flexible nature, the ligand-binding domain of a modified estrogen receptor (ER) was linked to the C-terminal of Oct-3/4 via a flexible linker containing GGGGS repeats (Fig. 1A) [21, 22]. In the absence of OHT, there is a high turnover of the fusion protein but binding to the OHT ligand stabilises the fusion protein, thereby providing a rapid inducible response [21]. We generated fusion constructs with flexible linkers containing 2, 4 and 6 repeats to better preserve Oct-3/4 functionality.

**Figure 1.**
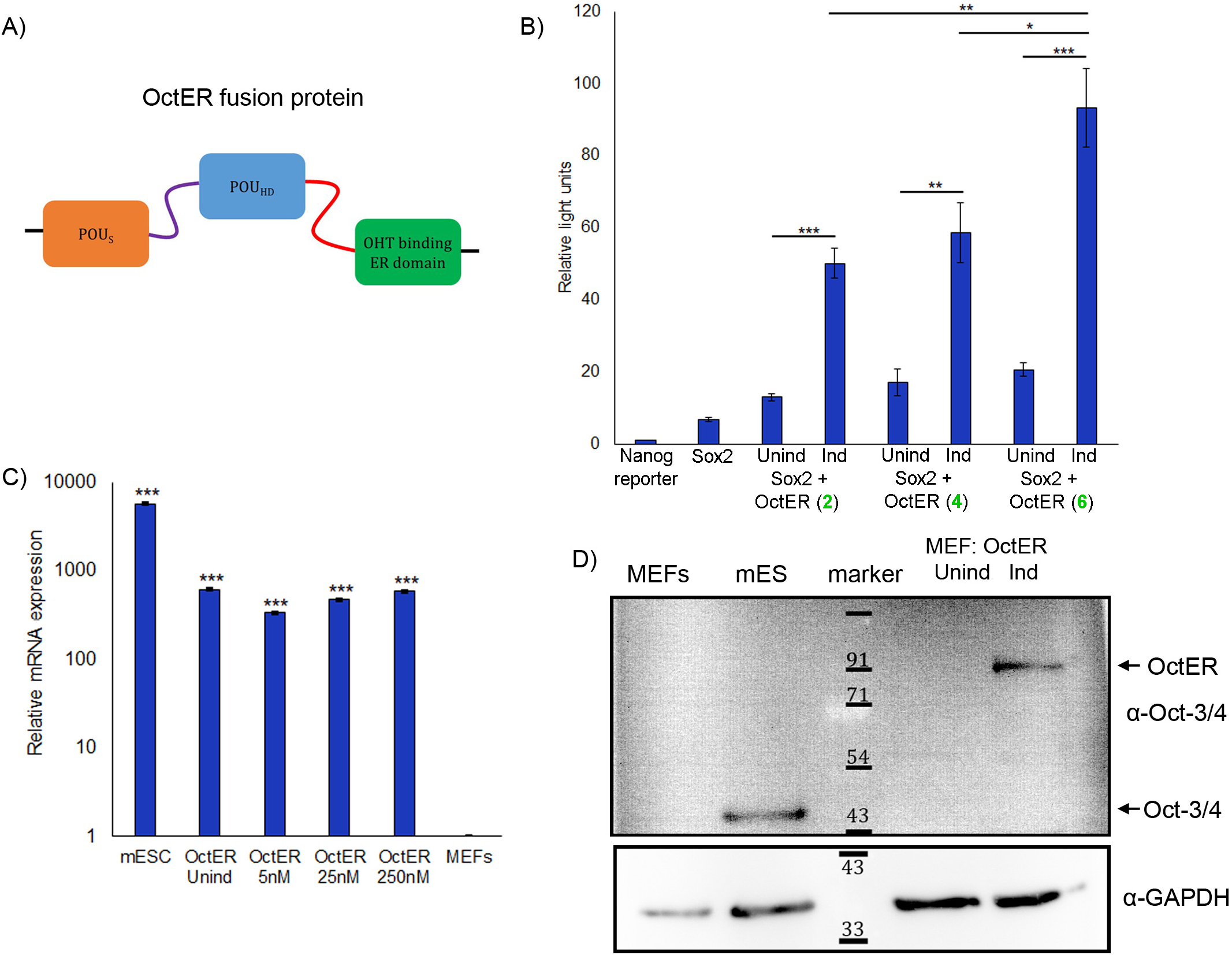
Constructing a tamoxifen-inducible Oct-3/4 to study changes in the early stages of reprogramming. A) Schematic of the OHT-inducible Oct-3/4 construct (OctER) showing the two DNA binding domains of Oct-3/4, POU_S_ (orange box) and POU_HD_ (blue box) connected by a native linker (purple line). The modified OHT binding domain (green box) is linked to Oct-3/4 via an introduced flexible linker (red line). B) A Nanog promoter luciferase reporter was used to determine the length of flexible linker required for optimal functioning of OctER along with Sox2, induced by 100 nM OHT. Reporter activity increased with increasing lengths of the linker suggesting greater separation between Oct-3/4 and modified ER aids transcriptional activity. Luciferase activity is normalized to basal Nanog promoter levels. Numbers in brackets signify number of linker repeats. Unind – Uninduced, Ind – Induced, N=4. C) Comparison of Oct-3/4 mRNA levels in mESC with MEFs expressing OctER under different doses of OHT. OctER mRNA levels are not altered by altered dose of OHT. N=3. D) Western blot of ectopic OctER protein levels in MEFs. MEFs do not natively express Oct-3/4. OctER was detected only in the presence of OHT (100 nM) and shows a higher molecular weight than endogenous Oct-3/4 due to addition of ER domain. Markers indicated in kD. Unind – Uninduced, Ind – Induced. Values represent mean + SEM, *P < 0.05, **P < 0.01, and ***P < 0.001

To test the function of these Oct-3/4-ER fusions, we used a Nanog promoter luciferase construct (Nanog-luc) with a functional Oct-3/4 and Sox2 binding site [23]. On its own, Nanog-luc was not activated by the synthetic ER ligand OHT (Fig. 1B), nor did co-expression of Sox2 show appreciable induction. However, co-expression of the different OctER fusions along with Sox2 showed an increase in Nanog promoter activity which was inducible by OHT (Fig. 1B). Notably, increased Nanog-luc expression corresponded to increased linker length (Fig. 1B). We conclude that greater separation between the modified ER and Oct-3/4 permitted better transcriptional function, and therefore used the inducible OctER with 6 flexible repeats for all further experiments.

In MEFs, ectopic OctER RNA levels were lower than Oct-3/4 levels in mESC and were unaffected by OHT (Fig. 1C). However, as expected, at the protein level, compared to endogenous Oct-3/4 protein (45 kD), the 80 kD OctER fusion protein was detectable only in OHT-induced MEFs and not in uninduced (Fig. 1D). Thus, the mRNA is stably expressed and the OctER fusion is inducible at the protein level, consistent with a more rapid response time compared to the TET system which requires transcriptional induction [24, 25].

### OctER activity is dose-dependent and rapid

To determine the induction characteristics of OctER protein, we analysed the levels of OctER in response to varying doses of OHT. 48 hours after transduction of MEFs with OctER lentivirus, increasing amounts of OHT (2.5 to 500 nM) were added. Following cell fractionation (Fig. S1), OctER protein was detectable in nuclear fractions after addition of OHT. The increase in levels of OctER in the nucleus correlated with the dose of OHT (Fig.2A, B), with no significant change in OctER RNA levels (Fig. 1C). Thus, OctER protein availability is dependent on OHT dose.

**Figure 2.**
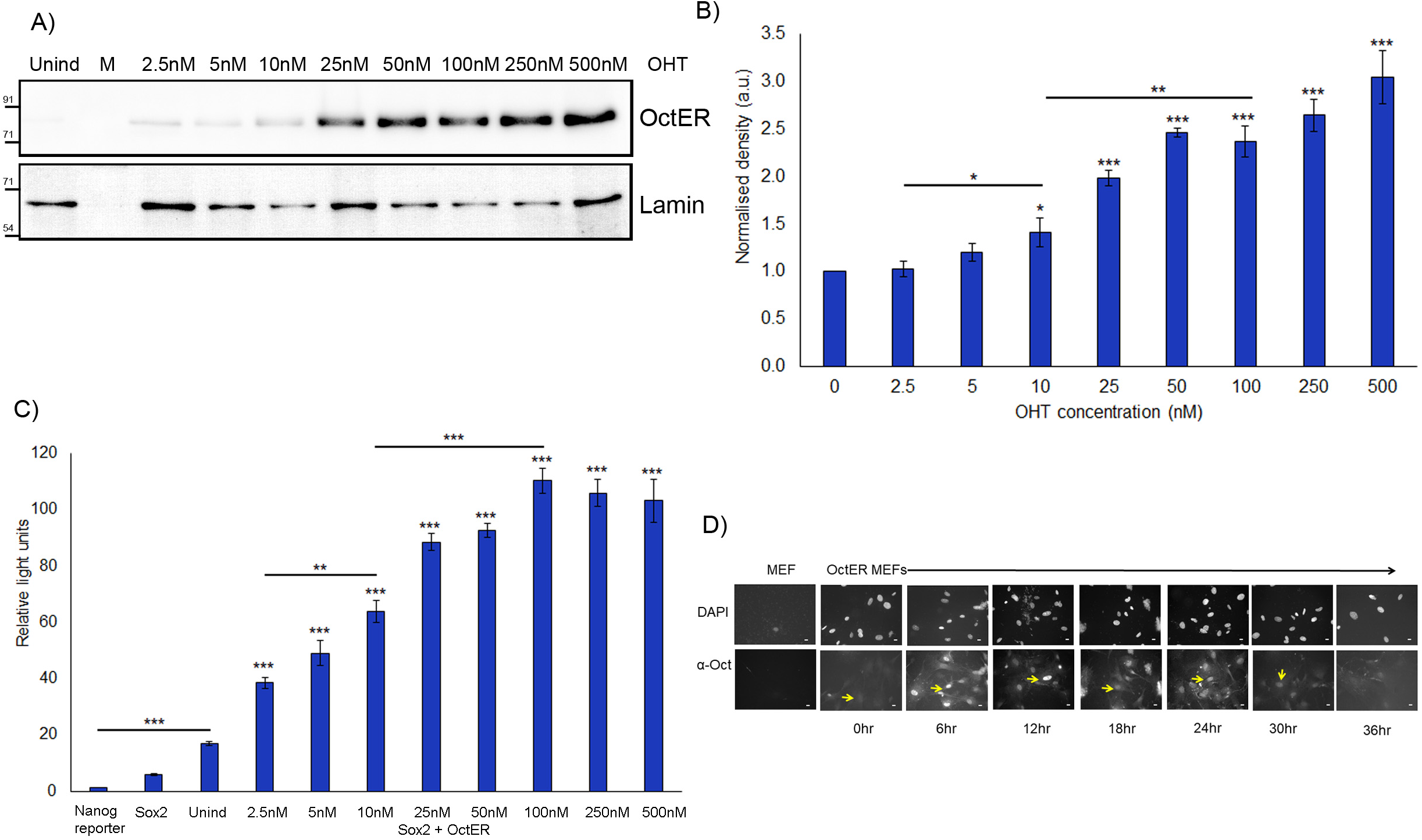
OctER activity is rapidly induced and dose-dependent. A) Western blots show that increasing doses of OHT lead to increasing levels of OctER in the nuclear fraction of OctER-expressing MEFs. Lamin B1 - loading control for nuclear fraction. Markers (M) indicated in kD. Unind – Uninduced B) Densitometric analysis of western blots show increased nuclear OctER with increasing doses of OHT. Values are normalised to Lamin B1 and plotted with respect to uninduced (0), N=4. C) Nanog promoter luciferase reporter activity reveals that OctER activity is OHT inducible. Reporter activity increased with increasing amounts of OHT showing that OctER activity is OHT dose dependent. Luciferase activities are shown relative to basal mouse Nanog promoter levels, N=4. Statistical analysis for induced samples were calculated with respect to uninduced. Unind – Uninduced D) Immunofluorescence staining for OctER in transduced MEF shows persistent nuclear localisation after a single pulse of 500 nM OHT. OctER is found in the nucleus by 6 hours post induction and persists up to 30 hours. Scale bar 20 μm. Values represent mean + SEM, *P < 0.05, **P < 0.01, and ***P < 0.001

To determine if OctER function was also OHT dose-dependent, we used the Nanog promoter luciferase assay. Co-expression of OctER with Sox2 led to a dose-dependent increase in Nanog promoter activity only when OHT was added (Fig. 2C). Nanog promoter activity was induced more than two fold by as low a dose as 2.5 nM OHT, plateauing at 100 nM with > 5 fold induction (Fig. 2C). Thus, OctER transcriptional activation of a canonical Oct-3/4 transcriptional target is OHT-inducible and can be modulated across a 40-fold dynamic range of ligand.

As complete reprogramming of MEFs requires 21 days, we determined the duration of OctER nuclear persistence. After a single pulse of 500 nM OHT, OctER was ‘chased’ in MEF nuclei for 48 hours. OctER was detected in MEF nuclei as early as 6 hours, and detectable for up to 30 hours (Fig. 2D). Thus, daily addition of OHT is sufficient to ensure that OctER is continuously available in MEF nuclei for reprogramming.

### OctER can reprogram MEFs

To test whether OctER can substitute for Oct-3/4 in reprogramming, we used MEFs isolated from OG-2 mice where a transgenic Oct-3/4 promoter-enhancer drives eGFP expression [26], enabling easy visual identification of reprogrammed cells. MEFs were transduced with a cocktail of either OctER, Klf4, Sox2, c-Myc (OctER + KSM) viruses or a positive control cocktail containing un-modified Oct-3/4, Klf4, Sox2, c-Myc (Oct-3/4 + KSM) (Fig. 3A), and treated with OHT daily. Colonies with compact morphology were apparent as early as 5 days post induction, in both positive control Oct-3/4 + KSM and test OctER + KSM experiments, and Oct-eGFP^+^ cells appeared within 10 days (Fig. 3B). Reprogramming efficiency of OctER + KSM and Oct-3/4 + KSM was calculated based on the number of compact alkaline phosphatase colonies seen at day 21 (Fig. 3C): the efficiency of OctER was lower than that of Oct-3/4, suggesting that despite efficient activation of the isolated Nanog promoter, composite functions required during reprogramming were less than wildtype Oct-3/4 (Fig. 3D). These experiments show that OctER can substitute for Oct-3/4 to reprogram MEFs into iPSCs, albeit at lower efficiency.

**Figure 3.**
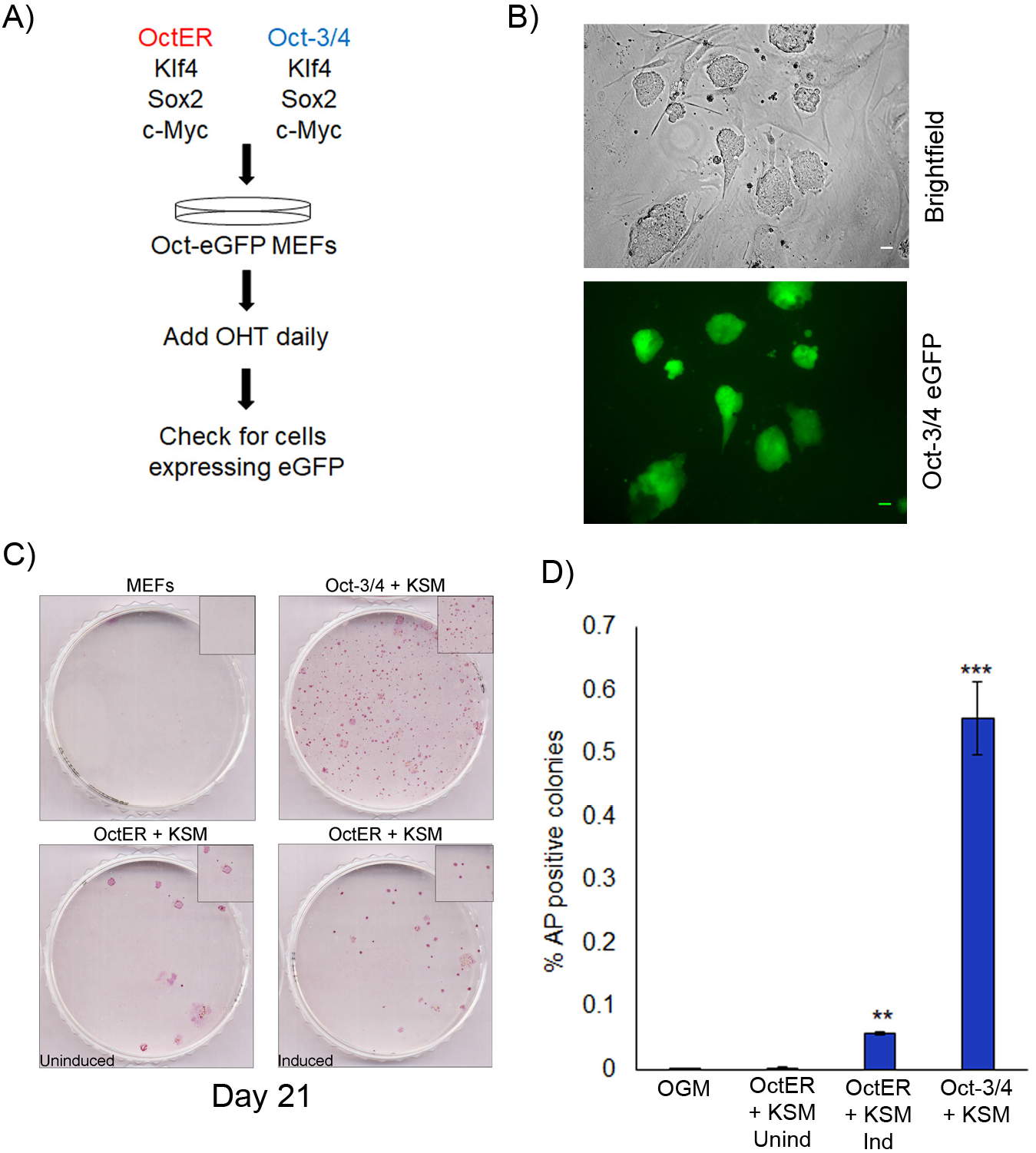
OctER can reprogram MEFs in combination with Klf4, Sox2 and c-Myc. A) Schematic used to reprogram MEFs to iPSC using either OctER + KSM or Oct-3/4 + KSM B) Reprogrammed OctER + KSM iPSC colonies express eGFP. Representative images: bright field (upper panel), and eGFP expressed from the transgenic Oct-3/4 promoter (lower panel). Scale bar 20 μm. C) iPSC colonies express alkaline phosphatase (AP). AP staining was used to determine the percentage of reprogrammed colonies: representative images of each condition are shown, insets display a magnified portion. Uninduced OctER sample shows dispersed cells while induction of OctER generates colonies resembling compact eGFP+ morphology of native Oct-3/4 expressing iPSC. D) Quantification of AP+ colonies produced using OctER + KSM vs. Oct-3/4 + KSM. Percentages were calculated based on the number of compact AP+ colonies obtained versus total number of cells plated, N=3. Values represent mean + SEM, *P < 0.05, **P < 0.01, and ***P < 0.001

### iPSC derived from OctER + KSM reprogramming are pluripotent

To determine if colonies with typical iPSC morphology obtained from OctER + KSM transduction were completely reprogramed, 4 colonies were picked and serially passaged for ten generations in the absence of OHT. All 4 colonies continued to express the transgenic Oct-eGFP reporter and were positive for alkaline phosphatase (Fig. 4A). mRNA levels for key pluripotency markers Cdh1, Dppa3, Nanog, Oct-3/4 (endogenous) and Sox2 in OctER + KSM iPSC were similar to levels seen in mESC (Fig. 4B). Simultaneously, OctER + KSM iPSC showed suppression (near absence) of key somatic markers, namely Cdh2, Snai1, Snai2, and Thy1 (Fig. 4C). OctER + KSM iPSC colonies also expressed pluripotency markers SSEA-1 and Nanog, further indicating OctER + KSM iPSC have successfully reprogrammed (Fig. 4D). Thus, iPSC derived from OctER-driven reprogramming achieved lasting OHT independence due to activation of endogenous Oct-3/4 and canonical pluripotency markers, and suppression of somatic cell markers.

**Figure 4.**
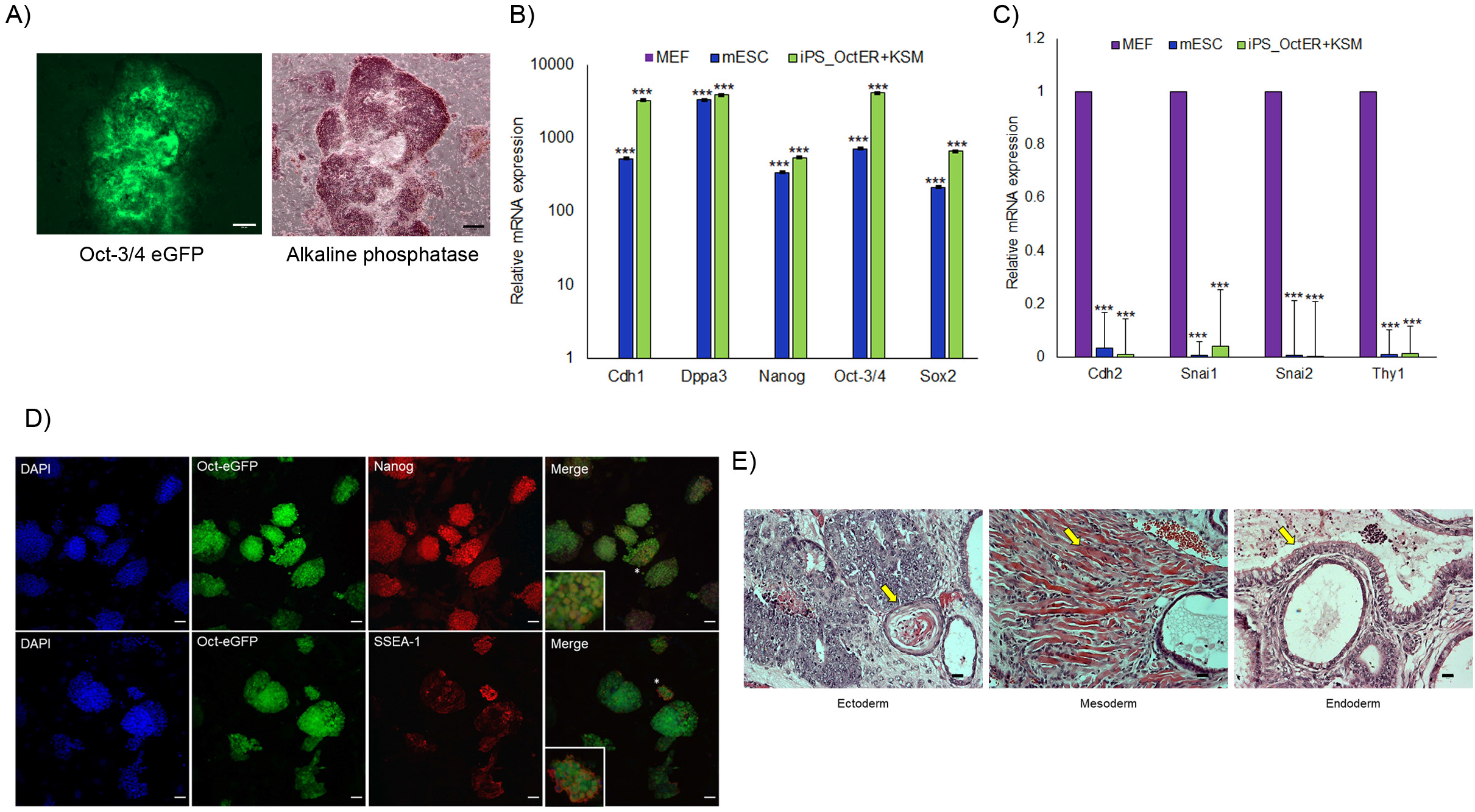
iPSC derived from OctER + KSM reprogramming are pluripotent. A) Representative image of an OctER + KSM iPSC colony after ten passages in the absence of inducer (OHT). The OctER + KSM iPSC colony is eGFP+ (left panel) and AP+ (right panel). Scale bar 200 μm. B) Quantification of mRNA levels of key pluripotency markers E-cadherin (Cdh1), Stella (Dppa3), Nanog, Oct-3/4, Sox2 in MEFs, mESC and OctER + KSM iPSC. Oct represents endogenous Oct-3/4 mRNA, N=3. C) Quantification of mRNA levels of key somatic markers N-cadherin (Cdh2), Snail (Snai1), Slug (Snai2), Thy1 in MEFs, mESC and OctER + KSM iPSC., N=3. D) Representative image of OctER + KSM iPSC colonies showing expression of Nanog and SSEA-1. Inset shows a single magnified colony (* in original image). Scale bar 50 μm E) H&E staining of sectioned teratoma derived from OctER + KSM iPSC. Germ layers were identified using distinguishing features (arrow). Keratinocyte rosette (ectoderm), striated muscle (mesoderm), ciliated columnar epithelium (endoderm). Scale bar 20 μm. Values represent mean + SEM, *P < 0.05, **P < 0.01, and ***P < 0.001

Further, OctER + KSM iPSC were able to form teratomas in nude mice. Histological analysis revealed the presence of tissue representing all 3 germ layers: ectoderm (keratinocyte rosette), endoderm (striated muscles) and mesoderm (ciliated columnar epithelia) (Fig. 4E). Thus, OctER can substitute for Oct-3/4 in reprogramming MEFs to a pluripotent state.

### Early effects of Klf4 and Oct-3/4 on MEF proliferation during reprogramming

The process of reprogramming while inefficient and stochastic, can be broadly classified into two major phases, initiation and maturation, where the first few days are associated with increased proliferative rates [9]. To determine the effects of Klf4 and Oct-3/4 on MEF proliferation, we focussed on the first 5 days of reprogramming. When expressed individually in MEFs, neither Klf4 or Oct-3/4 affected proliferation (Fig. 5A). However, when Klf4 and Oct-3/4 were expressed together, proliferation increased (Fig. 5A).

**Figure 5.**
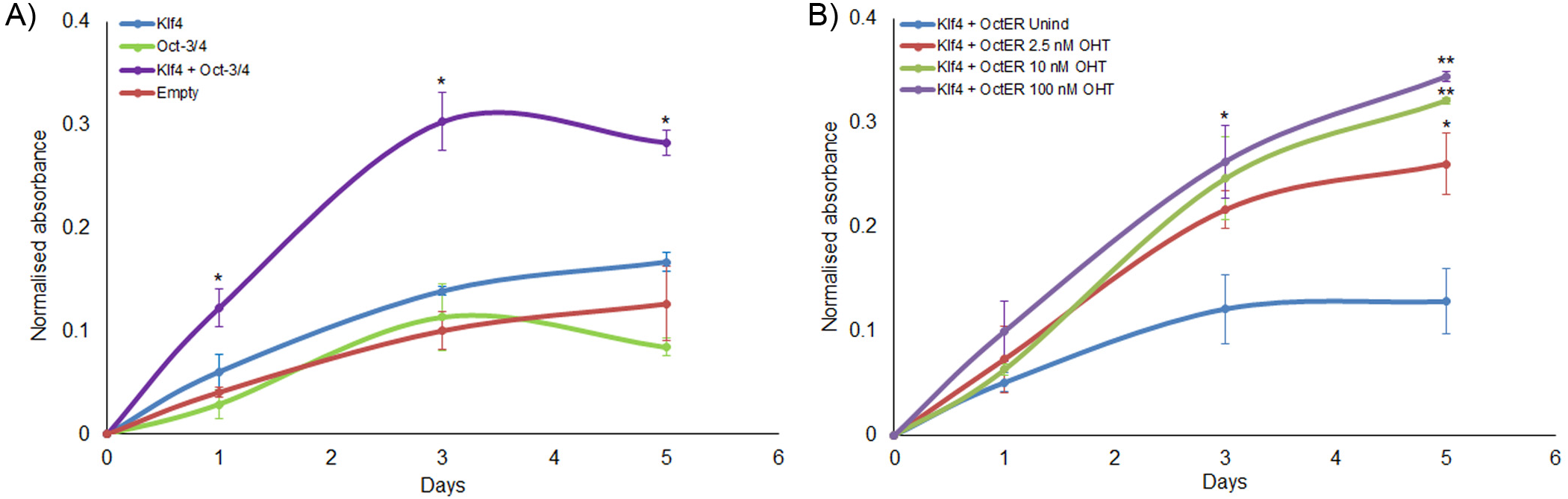
Early effects of Klf4 and Oct-3/4 on MEF proliferation during reprogramming. A) Klf4 and Oct-3/4 when expressed alone did not alter MEF proliferation, but when expressed together enhance MEF proliferation. Proliferation rates were compared to empty vector (Empty), measured using WST assay, N=3. B) Co-expression of OctER with Klf4 permits OHT-modulated MEF proliferation. In the uninduced state, MEF proliferation is unaffected. On addition of increasing amounts of OHT, there is a dose-dependent increase in MEF proliferation. Proliferation rates are compared to uninduced, N=3. Unind – Uninduced. Values represent mean + SEM, *P < 0.05, **P < 0.01, and ***P < 0.001

To determine if the changes in MEF proliferation by Klf4 and Oct-3/4 could be altered by a graded increase in Oct-3/4 levels, we used OctER induction. MEFs were transduced with Klf4 and OctER and subjected to dose-dependent induction of OctER function with 2.5 nM, 10 nM, 100 nM OHT (corresponding to 25%, 50% and 100% of Nanog promoter activation) (Fig. 2C). Without induction, Klf4 + OctER was similar to Klf4 alone, and did not change the slow proliferation (Fig. 5B). Addition of OHT caused an enhancement of proliferation that was dose-dependent, resulting in a graded increase in proliferation (Fig. 5B). Together, these results support the view that the effects of Klf4 and Oct-3/4 on MEF proliferation are synergistic and occur very early during reprogramming.

### Klf4 and Oct-3/4 target the Cyclin A2 promoter

To determine potential cell cycle targets of Klf4 and Oct-3/4, we mined published ChIP-seq data sets from NCBI GEO (Table S1) for links between Klf4 and Oct-3/4 and cell cycle genes (Table S2). Bioinformatic analysis showed that Oct-3/4 and Klf4 were enriched at the Cyclin A2 promoter in mESC and during early reprogramming (Fig. S2). Cyclin A2 is a key positive regulator of the cell cycle in mESC where it is highly expressed (Fig. S3) [1, 27], consistent with the hypothesis that induction of Cyclin A2 may be targeted by OctER + Klf4 to speed up the cell cycle and facilitate reprogramming. In-silico analysis of the Cyclin A2 promoter revealed potential Klf4 and Oct-3/4 motifs near the TSS [28, 29]. The Cyclin A2 promoter is under constant repression mediated by negative regulatory elements such as the cell cycle responsive element (CCRE) which is targeted by several epigenetic modifiers that control Cyclin A2 expression in somatic cells, as well as G1-S transcriptional regulators such as E2Fs and RB [30–34]. Positive transcriptional regulation of Cyclin A2 has been reported but comparatively less is known about this aspect [35]. To functionally assess binding of Klf4 and Oct-3/4 in the bioinformatically identified region, we used ChIP in mESC (Fig. 6A upper panel). Both endogenous Klf4 and Oct-3/4 showed enrichment at the predicted binding sites confirming our curation of the published ChIP-Seq analysis (Fig. 6A, middle and lower panels). Direct association of these transcription factors at the Cyclin A2 promoter is consistent with a potential regulatory role.

**Figure 6.**
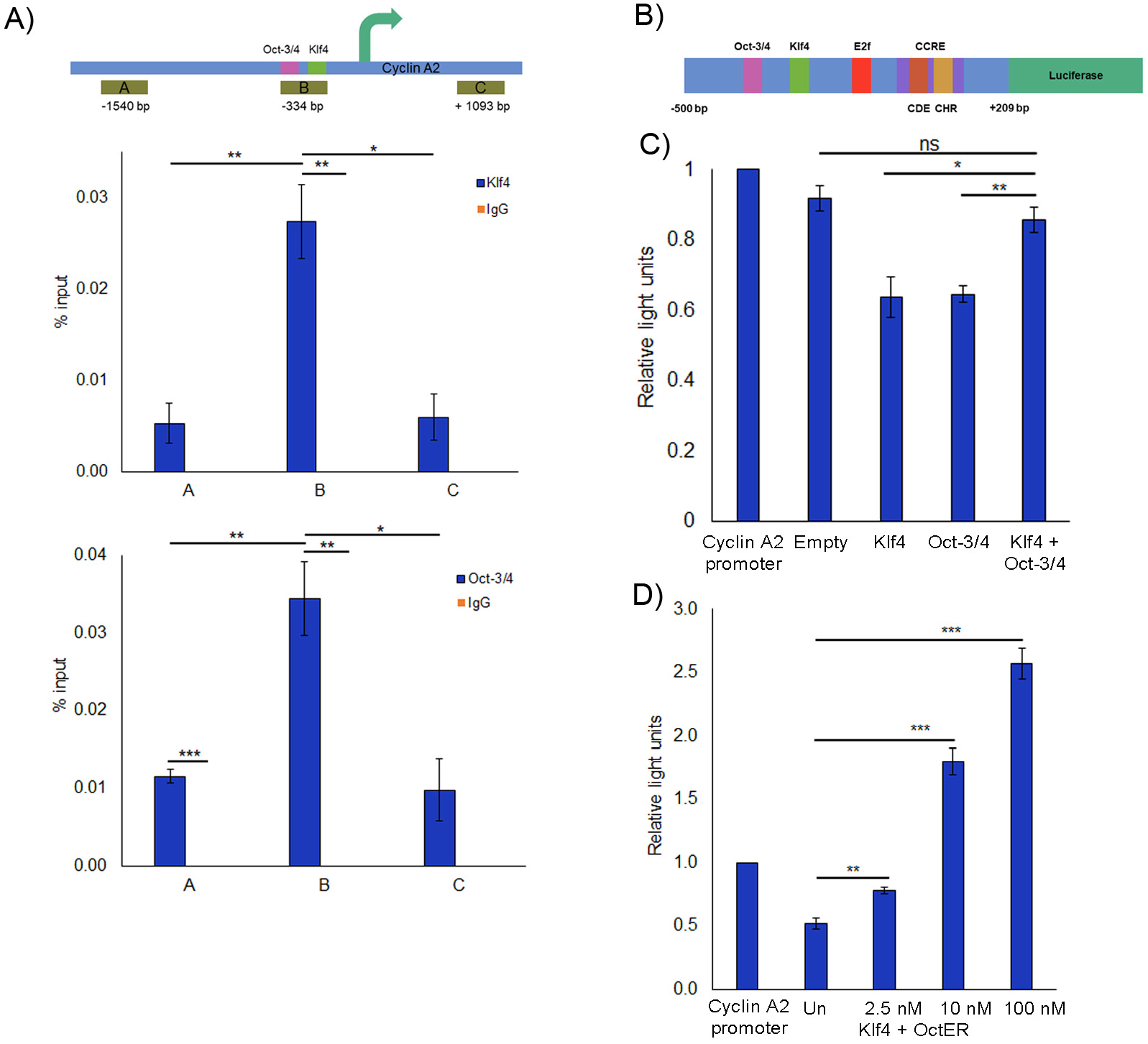
Oct-3/4 and Klf4 target the Cyclin A2 promoter. A) Upper panel: Schematic showing proximal promoter region of mouse Cyclin A2 genomic locus (arrow indicates TSS). Potential Klf4 and Oct-3/4 binding sites are marked in purple and green. Regions A (−1540 bp to −1385 bp), B (−336 bp to 154 bp) and C (1092 bp to 1243 bp) of the Cyclin A2 locus were tested for chromatin association of the reprogramming factors. Middle panel: ChIP analysis verifying enrichment of Klf4 at predicted binding site in Cyclin A2 promoter in mESC. Values are shown as % input along with IgG control, N=3. Lower panel: ChIP analysis verifying enrichment of Oct-3/4 at predicted binding site in Cyclin A2 promoter in mESC. Values are shown as % input along with IgG control, N=3. B) Schematic of mouse Cyclin A2 promoter region (−500 bp to +209 bp) cloned upstream of luciferase reporter. Known regulatory elements {E2F binding site, cell cycle responsive element (CCRE) comprising the cell cycle dependent element (CDE) and cell cycle gene homology region (CHR)} are marked. Sites predicted to bind Klf4 and Oct-3/4 are also shown. C) Effect of Klf4 and Oct-3/4 (alone or in combination) on Cyclin A2 promoter activity. Luciferase activity is normalised to basal level of Cyclin A2 reporter, N=3. D) OHT dose-dependent increase in Cyclin A2 promoter luciferase reporter activity under control of OctER when co-expressed with Klf4. Luciferase activity is normalised to basal level of Cyclin A2 reporter, N=3. Values represent mean + SEM, *P < 0.05, **P < 0.01, and ***P < 0.001

To verify if Klf4 and Oct-3/4 directly regulate Cyclin A2 expression, we cloned −500 bp to +209 bp of the mouse Cyclin A2 gene upstream of luciferase (Fig. 6B). This region includes the predicted Klf4 and Oct-3/4 binding sites verified in Fig. 6A, as well as the CCRE, and the first exon. In Cos7 cells, expression of either Klf4 or Oct-3/4 individually led to a slight reduction in Cyclin A2 promoter activity (Fig. 6C). However, co-expression of Klf4 and Oct-3/4 restored Cyclin A2 promoter activity (Fig. 6C).

To determine if the level of Oct-3/4 expression affects Cyclin A2 promoter activity in presence of Klf4, we used the inducible OctER. In absence of induction, OctER + Klf4 showed a reduction in Cyclin A2 promoter activity (Fig. 6D), as when Klf4 was expressed alone (Fig. 6C). However, on addition of low doses of OHT (2.5 nM, low OctER activity), the repression was reversed (Fig. 6D). In fact, higher doses of OHT were able to super-induce Cyclin A2 promoter activity (Fig. 6D). These findings suggest that there is a threshold of Oct-3/4 required to completely overcome repressive effects on the Cyclin A2 promoter, enabling high level expression of Cyclin A2. Taken together, this data suggests that Klf4 and Oct-3/4 work synergistically to reverse the repression associated with the Cyclin A2 proximal regulatory region.

### Klf4 and Oct-3/4 affect Cyclin A2 expression in MEFs

To determine whether ectopically expressed Klf4 and Oct-3/4 altered Cyclin A2 expression during early reprogramming in MEFs, we used the same experimental protocol as in Fig. 5. When Oct-3/4 or Klf4 were expressed individually in MEFs, there was no change in the levels of Cyclin A2 (Fig.7A). However, co-expression of Klf4 and Oct-3/4 induced a burst in Cyclin A2 mRNA levels (Fig. 7A).

**Figure 7.**
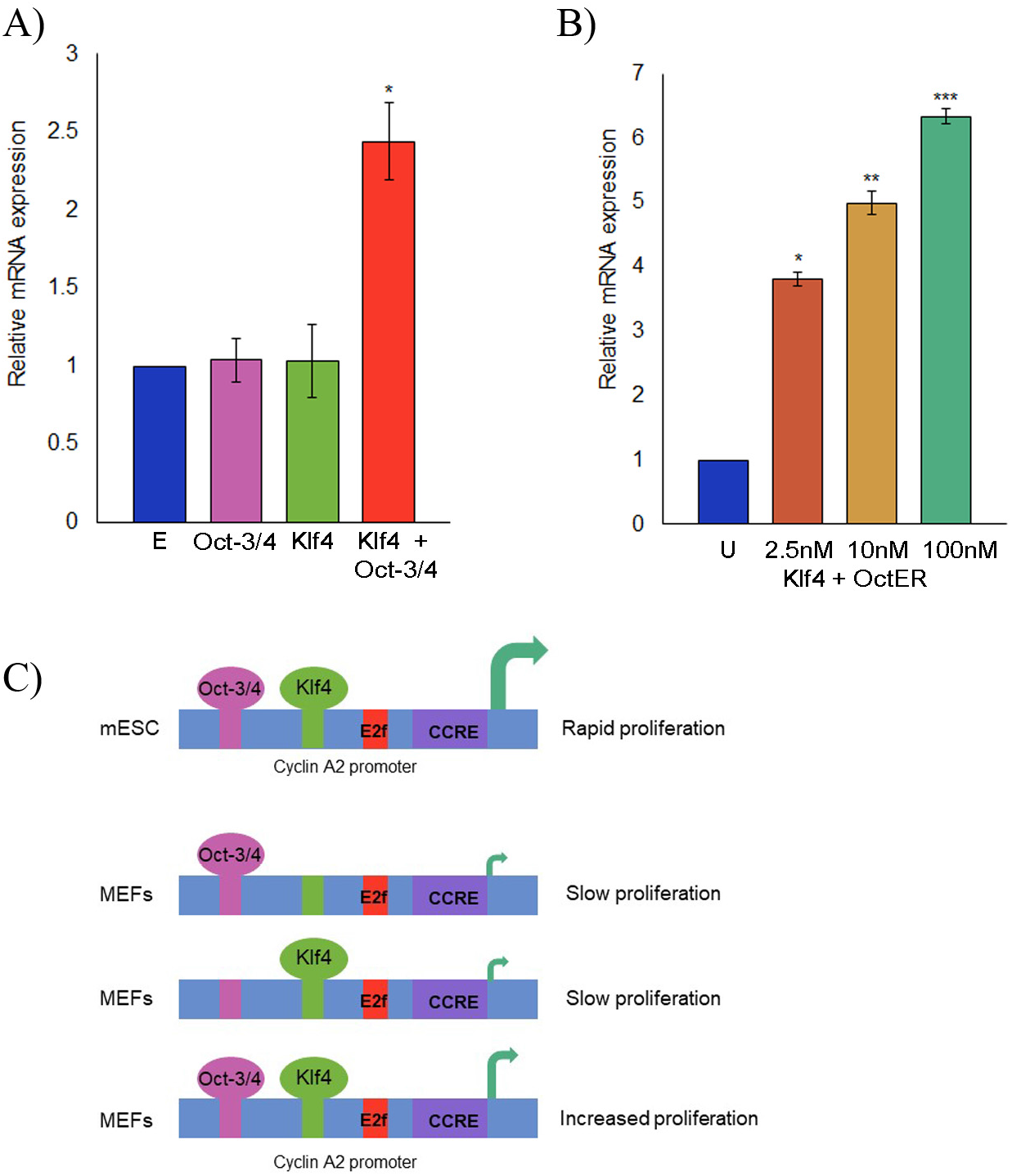
Oct-3/4 and Klf4 together activate Cyclin A2 expression in MEFs during early reprogramming. A) Cyclin A2 mRNA expression is unchanged in MEFs when Oct-3/4 or Klf4 are expressed individually, but is enhanced when the two factors are expressed together, N=3. E - Empty. B) OHT dose-dependent enhancement of Cyclin A2 mRNA expression in MEFs when OctER and Klf4 are co-expressed., N=3. U – Uninduced. C) Model illustrates the hypothesis that synergistic induction of Cyclin A2 promoter activity by Klf4 and Oct-3/4 in MEFs leads to an early enhancement of proliferation that is conductive to reprogramming. Values represent mean + SEM, *P < 0.05, **P < 0.01, and ***P < 0.001

To determine whether induction of Cyclin A2 was affected by the levels of Oct-3/4, OctER was co-expressed with Klf4 in MEFs and induced with OHT. Even at low doses of OHT, there was a substantial increase in Cyclin A2 mRNA levels, whereas at higher doses (corresponding to increased levels/function of OctER), Cyclin A2 expression was superinduced (Fig. 7B). Thus, the expression profile of Cyclin A2 transcript correlated well with the patterns of MEF proliferation observed (Fig. 5A), where individual factors could not reverse the repression of Cyclin A2 expression (Fig. 7B) and hence could not enhance proliferation (Fig.5A). However, when co-expressed, Klf4 and Oct-3/4 induced Cyclin A2 expression as well as increased MEF proliferation (Fig. 5A, 7A, 7B). Taken together, our results show that the early enhancement of the cell cycle during reprogramming may be mediated in part by Klf4 and Oct-3/4 synergistically targeting Cyclin A2 expression.

## Discussion

Cell cycle progression is mediated by a series of inter-connected regulatory loops whose control is still incompletely understood. A key question concerns the variable nature of cell cycle speed or duration in somatic vs. stem cells, primarily determined by the length of G1 [3]. In mESC, the rapid proliferative rate is attributed to the truncation of G1, which is thought to act as a protective mechanism against differentiation cues which act during this cell cycle stage [5, 36]. By comparison with somatic cells, a number of modifications to the cell cycle have been identified in mESC, including high/non oscillatory expression of positive cell cycle regulators, repression or inactivation of negative cell cycle regulators, and chromatin remodelling all of which contribute to rapid proliferation [1, 2, 5, 37].

By contrast to ESC, somatic cells have a longer G1 resulting in slower proliferation. In the context of reprogramming, selection of somatic cells with a shorter G1, using only rapidly proliferating cells, or overexpressing G1 Cyclin-Cdks such as Cyclin E and Cdk2, all lead to increased reprogramming efficiency [10–12]. Thus, cell cycle rate in particular the G1–S transition is a barrier to reprogramming. Since the path to pluripotency appears to require an accelerated cell cycle, somatic cell reprogramming is an excellent system for studying the impact of altered cell cycle duration.

Changes in the cell cycle occur during the early stages of reprogramming, when cells are still under the influence of exogenously expressed Yamanaka factors [9]. Modulating the levels of the Yamanaka factors also enhances reprogramming and may underlie the finding that only a small subset of cells that receive the appropriate ratios of pluripotency factors undergo successful reprogramming [38, 39]. To address the role of Klf4 and Oct-3/4 in early reprogramming, we created an inducible Oct-3/4 (OctER).

We show that OctER can replace Oct-3/4 in the Yamanaka quartet to reprogram MEFs to a pluripotent state. The lower efficiency of OctER may reflect insufficient linker length to prevent steric hindrance from the modified ER domain, which, being similar in size to that of Oct-3/4, might require greater separation than smaller fusion tags such as eGFP. While OctER efficiency is lower than Oct-3/4, it does produce fully functional iPSC in combination with Klf4, Sox2 and c-Myc: these iPSC are pluripotent as evidenced by RNA and protein analysis, and generate teratomas with all the three germ layers. Thus, the OctER construct satisfies the requirements for probing changes in the cell cycle along with Klf4.

Our study shows that when over-expressed in MEFs, Klf4 or Oct-3/4 individually do not alter the slow proliferation of these somatic cells. However, in combination Klf4 and Oct-3/4 together enhance MEF proliferation which could aid in reprogramming. Our finding is consistent with earlier reports that sequential addition of Klf4 and Oct-3/4, followed by c-Myc and finally Sox2 led to better reprogramming efficiency [38]. While previous studies have focussed on the mesenchymal-to-epithelial transition as a barrier to reprogramming, our studies suggest that enhancing the cell cycle may also assist in transcending this hurdle [38]. Indeed, expression analysis of partially reprogrammed cells showed many cell cycle genes were either upregulated or downregulated in comparison to mESC, suggesting that further altering cell cycle gene expression could move partially reprogrammed cells towards pluripotency [40].

Mining published ChIP-seq data sets (Table S1) revealed that the Cyclin A2 promoter has potential binding sites for Klf4 and Oct-3/4. Cyclin A2 is a positive cell cycle regulator responsible for the G1-S phase transition [41]. Interestingly, unlike Cyclin E, knock-out of Cyclin A2 does not affect MEF proliferation as drastically as in mESC [27]. Over-expression of Cyclin A2 accelerates entry into S phase [42, 43]. It is possible that enhancing expression of Cyclin A2 along with the Yamanaka factors might increase reprogramming efficiency.

In somatic cells, the Cyclin A2 promoter is largely under transcriptional repression, released only during the G1-S transition [44, 45]. This regulation is primarily mediated by the SWI/SNF chromatin remodeler that binds to the Cyclin A2 promoter [30, 31]. Depending on which ATPase subunit of SWI/SNF complex (Brg1 or Brm) binds, Cyclin A2 is expressed or repressed [30, 31]. In mESC, the esBAF SWI/SNF complex contains Brg1 and is important for maintaining proliferation [37], since it co-occupies promoters with pluripotency factors to regulate gene expression [46]. Our finding that Klf4 and Oct-3/4 bind in the vicinity of Brg1 on the Cyclin A2 promoter, is consistent with the view that co-occupancy of this promoter is important for its activity.

In MEFs, Brm knockout cells proliferate faster and exhibit higher Cyclin A2 expression [30]. Further, co-expression of Brg1 with Oct-3/4, Klf4 and Sox2 increases efficiency of reprogramming [47]. Our data shows that on addition of Klf4 and Oct-3/4, there is increased expression of Cyclin A2 as well as enhanced proliferation. Potentially, Klf4 and Oct-3/4 may enable replacement of Brm on the Cyclin A2 promoter by Brg1, thus increasing expression and promoting an ESC-like cell cycle.

Oct-3/4 is well established to play a key role in pluripotency by regulating factors such as Nanog, as well as by negating activity of lineage-determining genes such as Cdx2 [23, 48]. Oct-3/4 further acts by regulating the rapid pluripotent cell cycle via repression of negative regulators p21 and p53 [5]. We now report a new role for Oct-3/4 in maintaining pluripotency via enhanced expression of the positive cell cycle regulator, Cyclin A2. By binding to and enhancing promoter activity at the Cyclin A2 locus, Oct-3/4 and Klf4 maintain the rapid proliferation in ESC. Taken together, we propose that during reprogramming, Oct-3/4 and Klf4 bind to the Cyclin A2 promoter and together upregulate its expression thereby creating a privileged state for somatic cell conversion to pluripotency.

## Acknowledgements

We thank Drs. M. Inamdar and MM Panicker for useful discussions, Debaraya Saha and Dr. Gunjan Purohit for critical comments on the manuscript, Dr. M. Tada and Dr. K. Hasegawa for Nanog luciferase reporter, Dr. G. Evan for pBABE-cMycER and Dr. H. Scholer for Oct-eGFP mice. We are grateful to inStem for access to Animal House, Confocal Imaging and Flow Cytometry Facility; and CSIR-CCMB for access to advanced instrumentation and thank the staff of Animal House, Instrumentation group, Advanced Imaging Facility and Tissue Culture facility. L.Z. was supported by a CSIR senior research fellowship, inStem core funds, CCMB core funds. We are grateful to Drs. S. Rampalli, G. R. Chandak and I. Siddiqi for generously providing bridging support to L.Z. Animal work was partially supported by National Mouse Research Resource (NaMoR) grant (BT/PR5981/MED/31/181/2012; 2013-2016) from DBT to NCBS-inStem. This work was supported by Indo-Danish and Indo-Australian grants from DBT to J.D. as well as core funds from inStem and CSIR-CCMB.

## Author contributions

L.Z. and J.D. designed the experiments. L.Z. performed the experiments. L.Z. and J.D. wrote the manuscript.

## Competing Interests

The authors declare no competing interests.

## Figure legends

## Notes

### Competing Interest Statement

The authors have declared no competing interest.

## References

[1] E. Stead, J. White, R. Faast, S. Conn, S. Goldstone, J. Rathjen, U. Dhingra, P. Rathjen, D. Walker, S. Dalton, Pluripotent cell division cycles are driven by ectopic Cdk2, cyclin A/E and E2F activities, Oncogene, 21 (2002) 8320–8333.

[2] H. Fujii-Yamamoto, J.M. Kim, K. Arai, H. Masai, Cell cycle and developmental regulations of replication factors in mouse embryonic stem cells, The Journal of biological chemistry, 280 (2005) 12976–12987.

[3] C. Lange, F. Calegari, Cdks and cyclins link G1 length and differentiation of embryonic, neural and hematopoietic stem cells, Cell cycle, 9 (2010) 1893–1900.

[4] M.J. Evans, M.H. Kaufman, Establishment in culture of pluripotential cells from mouse embryos, Nature, 292 (1981) 154–156.

[5] L. Zaveri, J. Dhawan, Cycling to Meet Fate: Connecting Pluripotency to the Cell Cycle, Front Cell Dev Biol, 6 (2018) 57.

[6] M.H.L. Snow, Gastrulation in the mouse: Growth and regionalization of the epiblast, Journal of embryology and experimental morphology, 42 (1977) 293–303.

[7] P. Savatier, S. Huang, L. Szekely, K.G. Wiman, J. Samarut, Contrasting patterns of retinoblastoma protein expression in mouse embryonic stem cells and embryonic fibroblasts, Oncogene, 9 (1994) 809–818.

[8] K. Takahashi, S. Yamanaka, Induction of pluripotent stem cells from mouse embryonic and adult fibroblast cultures by defined factors, Cell, 126 (2006) 663–676.

[9] J.M. Polo, E. Anderssen, R.M. Walsh, B.A. Schwarz, C.M. Nefzger, S.M. Lim, M. Borkent, E. Apostolou, S. Alaei, J. Cloutier, O. Bar-Nur, S. Cheloufi, M. Stadtfeld, M.E. Figueroa, D. Robinton, S. Natesan, A. Melnick, J. Zhu, S. Ramaswamy, K. Hochedlinger, A molecular roadmap of reprogramming somatic cells into iPS cells, Cell, 151 (2012) 1617–1632.

[10] S. Ruiz, A.D. Panopoulos, A. Herrerias, K.D. Bissig, M. Lutz, W.T. Berggren, I.M. Verma, J.C. Izpisua Belmonte, A high proliferation rate is required for cell reprogramming and maintenance of human embryonic stem cell identity, Current biology: CB, 21 (2011) 45–52.

[11] S. Guo, X. Zi, V.P. Schulz, J. Cheng, M. Zhong, S.H. Koochaki, C.M. Megyola, X. Pan, K. Heydari, S.M. Weissman, P.G. Gallagher, D.S. Krause, R. Fan, J. Lu, Nonstochastic reprogramming from a privileged somatic cell state, Cell, 156 (2014) 649–662.

[12] M. Roccio, D. Schmitter, M. Knobloch, Y. Okawa, D. Sage, M.P. Lutolf, Predicting stem cell fate changes by differential cell cycle progression patterns, Development, 140 (2013) 459–470.

[13] M. Nakagawa, M. Koyanagi, K. Tanabe, K. Takahashi, T. Ichisaka, T. Aoi, K. Okita, Y. Mochiduki, N. Takizawa, S. Yamanaka, Generation of induced pluripotent stem cells without Myc from mouse and human fibroblasts, Nature biotechnology, 26 (2008) 101–106.

[14] Z. Lin, P. Perez, D. Lei, J. Xu, X. Gao, J. Bao, Two-phase analysis of molecular pathways underlying induced pluripotent stem cell induction, Stem cells, 29 (2011) 1963–1974.

[15] Y. Buganim, D.A. Faddah, A.W. Cheng, E. Itskovich, S. Markoulaki, K. Ganz, S.L. Klemm, A. van Oudenaarden, R. Jaenisch, Single-cell expression analyses during cellular reprogramming reveal an early stochastic and a late hierarchic phase, Cell, 150 (2012) 1209–1222.

[16] M. Wernig, A. Meissner, J.P. Cassady, R. Jaenisch, c-Myc is dispensable for direct reprogramming of mouse fibroblasts, Cell stem cell, 2 (2008) 10–12.

[17] J. Jiang, Y.S. Chan, Y.H. Loh, J. Cai, G.Q. Tong, C.A. Lim, P. Robson, S. Zhong, H.H. Ng, A core Klf circuitry regulates self-renewal of embryonic stem cells, Nature cell biology, 10 (2008) 353–360.

[18] B.D. Rowland, D.S. Peeper, KLF4, p21 and context-dependent opposing forces in cancer, Nature reviews. Cancer, 6 (2006) 11–23.

[19] H. Niwa, J. Miyazaki, A.G. Smith, Quantitative expression of Oct-3/4 defines differentiation, dedifferentiation or self-renewal of ES cells, Nature genetics, 24 (2000) 372–376.

[20] S. Jerabek, F. Merino, H.R. Scholer, V. Cojocaru, OCT4: dynamic DNA binding pioneers stem cell pluripotency, Biochimica et biophysica acta, 1839 (2014) 138–154.

[21] T.D. Littlewood, D.C. Hancock, P.S. Danielian, M.G. Parker, G.I. Evan, A modified oestrogen receptor ligand-binding domain as an improved switch for the regulation of heterologous proteins, Nucleic acids research, 23 (1995) 1686–1690.

[22] X. Chen, J.L. Zaro, W.C. Shen, Fusion protein linkers: property, design and functionality, Advanced drug delivery reviews, 65 (2013) 1357–1369.

[23] T. Kuroda, M. Tada, H. Kubota, H. Kimura, S.Y. Hatano, H. Suemori, N. Nakatsuji, T. Tada, Octamer and Sox elements are required for transcriptional cis regulation of Nanog gene expression, Mol Cell Biol, 25 (2005) 2475–2485.

[24] A.M. Kringstein, F.M. Rossi, A. Hofmann, H.M. Blau, Graded transcriptional response to different concentrations of a single transactivator, Proceedings of the National Academy of Sciences of the United States of America, 95 (1998) 13670–13675.

[25] A.D. Kohn, A. Barthel, K.S. Kovacina, A. Boge, B. Wallach, S.A. Summers, M.J. Birnbaum, P.H. Scott, J.C. Lawrence, Jr., R.A. Roth, Construction and characterization of a conditionally active version of the serine/threonine kinase Akt, The Journal of biological chemistry, 273 (1998) 11937–11943.

[26] P.E. Szabo, K. Hubner, H. Scholer, J.R. Mann, Allele-specific expression of imprinted genes in mouse migratory primordial germ cells, Mechanisms of development, 115 (2002) 157–160.

[27] I. Kalaszczynska, Y. Geng, T. Iino, S. Mizuno, Y. Choi, I. Kondratiuk, D.P. Silver, D.J. Wolgemuth, K. Akashi, P. Sicinski, Cyclin A is redundant in fibroblasts but essential in hematopoietic and embryonic stem cells, Cell, 138 (2009) 352–365.

[28] I.V. Kulakovskiy, I.E. Vorontsov, I.S. Yevshin, R.N. Sharipov, A.D. Fedorova, E.I. Rumynskiy, Y.A. Medvedeva, A. Magana-Mora, V.B. Bajic, D.A. Papatsenko, F.A. Kolpakov, V.J. Makeev, HOCOMOCO: towards a complete collection of transcription factor binding models for human and mouse via large-scale ChIP-Seq analysis, Nucleic acids research, 46 (2018) D252–D259.

[29] O. Fornes, J.A. Castro-Mondragon, A. Khan, R. van der Lee, X. Zhang, P.A. Richmond, B.P. Modi, S. Correard, M. Gheorghe, D. Baranasic, W. Santana-Garcia, G. Tan, J. Cheneby, B. Ballester, F. Parcy, A. Sandelin, B. Lenhard, W.W. Wasserman, A. Mathelier, JASPAR 2020: update of the open-access database of transcription factor binding profiles, Nucleic acids research, 48 (2020) D87–D92.

[30] M. Coisy, V. Roure, M. Ribot, A. Philips, C. Muchardt, J.M. Blanchard, J.C. Dantonel, Cyclin A repression in quiescent cells is associated with chromatin remodeling of its promoter and requires Brahma/SNF2alpha, Molecular cell, 15 (2004) 43–56.

[31] S. Kadam, B.M. Emerson, Transcriptional specificity of human SWI/SNF BRG1 and BRM chromatin remodeling complexes, Molecular cell, 11 (2003) 377–389.

[32] J. Zwicker, F.C. Lucibello, L.A. Wolfraim, C. Gross, M. Truss, K. Engeland, R. Muller, Cell-Cycle Regulation of the Cyclin-a, Cdc25c and Cdc2 Genes Is Based on a Common Mechanism of Transcriptional Repression, Embo Journal, 14 (1995) 4514–4522.

[33] K.E. Knudsen, A.F. Fribourg, M.W. Strobeck, J.M. Blanchard, E.S. Knudsen, Cyclin A is a functional target of retinoblastoma tumor suppressor protein-mediated cell cycle arrest, The Journal of biological chemistry, 274 (1999) 27632–27641.

[34] A. Schulze, K. Zerfass, D. Spitkovsky, S. Middendorp, J. Berges, K. Helin, P. Jansen-Durr, B. Henglein, Cell cycle regulation of the cyclin A gene promoter is mediated by a variant E2F site, Proceedings of the National Academy of Sciences of the United States of America, 92 (1995) 11264–11268.

[35] M. Katabami, H. Donninger, F. Hommura, V.D. Leaner, I. Kinoshita, J.F. Chick, M.J. Birrer, Cyclin A is a c-Jun target gene and is necessary for c-Jun-induced anchorage-independent growth in RAT1a cells, The Journal of biological chemistry, 280 (2005) 16728–16738.

[36] D. Coronado, M. Godet, P.Y. Bourillot, Y. Tapponnier, A. Bernat, M. Petit, M. Afanassieff, S. Markossian, A. Malashicheva, R. Iacone, K. Anastassiadis, P. Savatier, A short G1 phase is an intrinsic determinant of naive embryonic stem cell pluripotency, Stem cell research, 10 (2013) 118–131.

[37] L. Ho, J.L. Ronan, J. Wu, B.T. Staahl, L. Chen, A. Kuo, J. Lessard, A.I. Nesvizhskii, J. Ranish, G.R. Crabtree, An embryonic stem cell chromatin remodeling complex, esBAF, is essential for embryonic stem cell self-renewal and pluripotency, Proceedings of the National Academy of Sciences of the United States of America, 106 (2009) 5181–5186.

[38] X. Liu, H. Sun, J. Qi, L. Wang, S. He, J. Liu, C. Feng, C. Chen, W. Li, Y. Guo, D. Qin, G. Pan, J. Chen, D. Pei, H. Zheng, Sequential introduction of reprogramming factors reveals a time-sensitive requirement for individual factors and a sequential EMT-MET mechanism for optimal reprogramming, Nature cell biology, 15 (2013) 829–838.

[39] B.W. Carey, S. Markoulaki, J.H. Hanna, D.A. Faddah, Y. Buganim, J. Kim, K. Ganz, E.J. Steine, J.P. Cassady, M.P. Creyghton, G.G. Welstead, Q. Gao, R. Jaenisch, Reprogramming factor stoichiometry influences the epigenetic state and biological properties of induced pluripotent stem cells, Cell stem cell, 9 (2011) 588–598.

[40] C. Chronis, P. Fiziev, B. Papp, S. Butz, G. Bonora, S. Sabri, J. Ernst, K. Plath, Cooperative Binding of Transcription Factors Orchestrates Reprogramming, Cell, 168 (2017) 442–459 e420.

[41] F. Girard, U. Strausfeld, A. Fernandez, N.J.C. Lamb, Cyclin-a Is Required for the Onset of DNA-Replication in Mammalian Fibroblasts, Cell, 67 (1991) 1169–1179.

[42] D. Resnitzky, L. Hengst, S.I. Reed, Cyclin A-associated kinase activity is rate limiting for entrance into S phase and is negatively regulated in G1 by p27Kip1, Mol Cell Biol, 15 (1995) 4347–4352.

[43] A.R. Rosenberg, F. Zindy, F. Ledeist, H. Mouly, P. Metezeau, C. Brechot, E. Lamas, Overexpression of Human Cyclin-a Advances Entry into S-Phase, Oncogene, 10 (1995) 1501–1509.

[44] X. Huet, J. Rech, A. Plet, A. Vie, J.M. Blanchard, Cyclin A expression is under negative transcriptional control during the cell cycle, Mol Cell Biol, 16 (1996) 3789–3798.

[45] B. Henglein, X. Chenivesse, J. Wang, D. Eick, C. Brechot, Structure and cell cycle-regulated transcription of the human cyclin A gene, Proceedings of the National Academy of Sciences of the United States of America, 91 (1994) 5490–5494.

[46] L. Ho, R. Jothi, J.L. Ronan, K. Cui, K. Zhao, G.R. Crabtree, An embryonic stem cell chromatin remodeling complex, esBAF, is an essential component of the core pluripotency transcriptional network, Proceedings of the National Academy of Sciences of the United States of America, 106 (2009) 5187–5191.

[47] N. Singhal, J. Graumann, G. Wu, M.J. Arauzo-Bravo, D.W. Han, B. Greber, L. Gentile, M. Mann, H.R. Scholer, Chromatin-Remodeling Components of the BAF Complex Facilitate Reprogramming, Cell, 141 (2010) 943–955.

[48] H. Niwa, Y. Toyooka, D. Shimosato, D. Strumpf, K. Takahashi, R. Yagi, J. Rossant, Interaction between Oct3/4 and Cdx2 determines trophectoderm differentiation, Cell, 123 (2005) 917–929.

[49] K. Suzuki, P. Bose, R.Y. Leong-Quong, D.J. Fujita, K. Riabowol, REAP: A two minute cell fractionation method, BMC Res Notes, 3 (2010) 294.

