## Supplementary Information for "Inducible expression of Oct-3/4 reveals synergy with Klf4 in targeting Cyclin A2 to enhance proliferation during early reprogramming"

Zaveri and Dhawan - Supplementary Information

**Supplementary figures**

Figure S1

Figure S2

Figure S3

**Supplementary tables**

Table S1: List of published ChIP-Seq data sets analysed containing keywords related to cell cycle, pluripotency, embryonic stem cells and reprogramming

Table S2: List of cell cycle genes analysed

Table S3: List of cloning primers

Table S4: List of plasmids used

Table S5: List of qRT-PCR primers used in this study for mRNA expression analysis

Table S6: List of ChIP primers

Table S7: List of Antibodies used

### Figure S1

A)

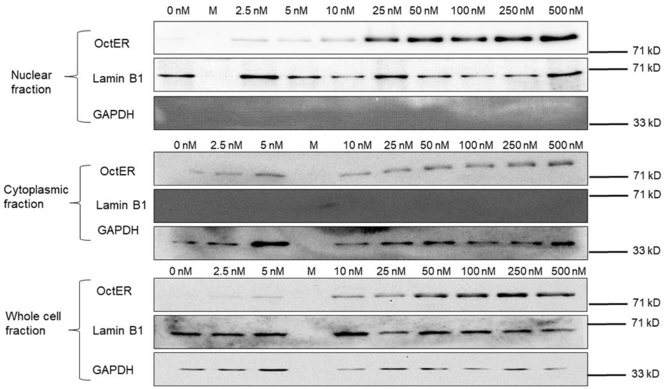
0 nM
M
2.5 nM
5 nM
10 nM
25 nM
50 nM
100 nM
250 nM
500 nM
Nuclear fraction
OctER
Lamin B1
GAPDH
71 kD
71 kD
33 kD
Cytoplasmic fraction
OctER
Lamin B1
GAPDH
71 kD
71 kD
33 kD
Whole cell fraction
OctER
Lamin B1
GAPDH
71 kD
71 kD
33 kD

### Figure S2

A)

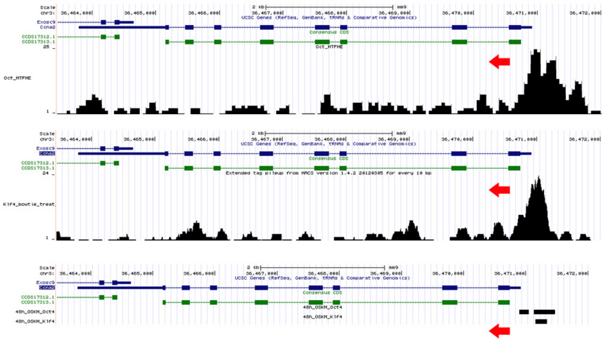
Scale
chr3:
2 kb
mm9
36,464,000
36,465,000
36,466,000
36,467,000
36,468,000
36,469,000
36,470,000
36,471,000
36,472,000
UCSC Genes (RefSeq, GenBank, tRNAs & Comparative Genomics)
Exosc9
Ccna2
Consensus CDS
CCDS17312.1
CCDS17313.1
25
Oct_MTFME
1
24
Extended tag pileup from MACS version 1.4.2 20120305 for every 10 bp
Klf4_bowtie_treat
48h_OSKM_Oct4
48h_OSKM_Klf4

Figure S3

A)

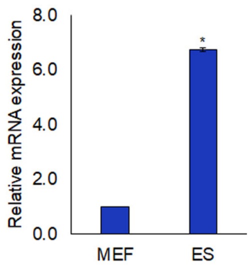

**Supplementary figure legends**

Figure S1

Western blots show that increasing doses of OHT lead to increasing levels of OctER in the nuclear fraction of OctER-expressing MEFs. Lamin B1 - loading control for nuclear fraction, GAPDH for cytoplasmic fraction. Markers (M) indicated in kD. Unind – Uninduced

Figure S2

Representative images from bioinformatic analysis of Cyclin A2 gene for enrichment of Oct-3/4 and Klf4 in mESC and MEFs during early reprogramming. Upper panel shows enrichment peak for Oct-3/4 (GSE44288), middle panel for Klf4 (GSE49848) and lower panel for both Oct-3/4 and Klf4 during early reprogramming (GSE90895). Red arrow indicates direction of transcription.

Figure S3

Comparison of mRNA levels of Cyclin A2 in mESC with MEFs. N=3.

Values represent mean + SEM, *P < 0.05, **P < 0.01, and ***P < 0.001

**Supplementary tables**

Table S1: List of published ChIP-Seq data sets analysed containing keywords related to cell cycle, pluripotency, embryonic stem cells and reprogramming

| GEO accession number | Transcription factor | Publication title |
|---|---|---|
| GSE52397 | Klf4 | C/EBPα poises B cells for rapid reprogramming into induced pluripotent stem cells |
| GSE90895 | Oct-3/4, Klf4 | Cooperative Binding of Transcription Factors Orchestrates Reprogramming |

| | | |
|---|---|---|
| GSE11724 | Oct-3/4 | Connecting microRNA genes to the core transcriptional regulatory circuitry of embryonic stem cells |
| GSE43231 | Oct-3/4 | Distinct and combinatorial functions of Jmjd2b/Kdm4b and Jmjd2c/Kdm4c in mouse embryonic stem cell identity |
| GSE36570 | Oct-3/4, Klf4 | Facilitators and impediments of the pluripotency reprogramming factors' initial engagement with the genome |
| GSE56312 | Oct-3/4 | Ground State Conditions Induce Rapid Reorganization of Core Pluripotency Factor Binding before Global Epigenetic Reprogramming |
| GSE67520 | Oct-3/4 | Hierarchical Oct4 Binding in Concert with Primed Epigenetic Rearrangements during Somatic Cell Reprogramming |
| GSE49848 | Klf4 | Klf4 and Klf5 differentially inhibit mesoderm and endoderm differentiation in embryonic stem cells |
| GSE44288 | Oct-3/4 | Master transcription factors and mediator establish super-enhancers at key cell identity genes |
| GSE74636 | Oct-3/4 | ChIP-seq analysis of genomic binding regions of five major transcription factors highlights a central role for ZIC2 in the mouse epiblast stem cell gene regulatory network. |
| GSE43275 | Oct-3/4 | Oct4 switches partnering from Sox2 to Sox17 to reinterpret the enhancer code and specify endoderm |
| GSE56138 | Oct-3/4 | Reorganization of enhancer patterns in transition from naive to primed pluripotency |
| GSE61475 | Oct-3/4 | Transcription factor binding dynamics during human ES cell differentiation |
| GSE46130 | Oct-3/4 | Transcriptional and epigenetic dynamics during specification of human embryonic stem cells |
| GSE92846 | Oct-3/4, Klf4 | Widespread Mitotic Bookmarking by Histone Marks and Transcription Factors in Pluripotent Stem Cells. |
| GSE22934 | Oct-3/4 | Wdr5 mediates self-renewal and reprogramming via the embryonic stem cell core transcriptional network |

Table S2: List of cell cycle genes analysed across ChIP-seq data sets for potential enrichment sites across the gene body and promoter region, allowing us to select Cyclin A2 as a potential direct target of Klf4 and Oct-3/4 during early reprogramming.

| Cell cycle gene |
|---|
| Cyclin A2 |
| Cyclin B |
| Cyclin D1 |
| Cyclin D2 |
| Cyclin D3 |
| Cyclin E |
| Cdk1 |
| Cdk2 |
| Cdk4 |
| Cdk6 |
| p21 (Cdkn1a) |
| p27 (Cdkn1b) |

Table S3: List of cloning primers

| Primer | Sequence (5' to 3') |
|---|---|
| AgeI-Koz-Oct Fwd | ACC GGT CCA CCT TCC GCC ACC ATG GCT GGA C |
| Oct3/4-NheI-EcoRI-Rev | GAA TTC CTC TCG CTA GCG TTT GAA TGC ATG GGA GAG C |
| PacI-CMV-Fwd | TTA ATT AAA AGC TTG GGA GTT CCG C |
| CMV-BamHI-Rev | GGA TCC TCT AGT AGA GTC GGT GTC |
| NheI-Oct3/4-(2 repeats)-ER-Fwd | GCT AGC GGA GGA GGA GGA TCA GGA GGA GGA GGA TCA CGA AAT GAA ATG GGT GCT TC |
| NheI-Oct3/4-(3 repeats)-ER-Fwd | GCT AGC GGA GGA GGA GGA TCA GGA GGA GGA GGA TCA GGA GGA GGA GGA TCA CGA AAT GAA ATG GGT GCT TC |

| | |
|---|---|
| NheI-Oct3/4-(4 repeats)-ER-Fwd | GCT AGC GGA GGA GGA GGA TCA GGA GGA GGA GGA TCA GGA GGA GGA GGA TCA GGA GGA GGA GGA TCA CGA AAT GAA ATG GGT GCT TC |
| NheI-Oct3/4-(6 repeats)-ER-Fwd | GCT AGC GGA GGA GGA GGA TCA GGA GGA GGA GGA TCA GGA GGA GGA GGA TCA GGA GGA GGA GGA TCA GGA GGA GGA GGA TCA CGA AAT GAA ATG GGT GCT TC |
| EcoRI-STOP-ER-Rev | GAA TTC ATC CGA TCG TGT TGG GGA AGC |
| NheI-CCNA2-5Pos-Fwd | GCT AGC TTG GGA CAG CAT TAT GAG ACC |
| CCNA2-TSS-Rev-HindIII | AAG CTT GAC CCG AAT GCC TCG AGG TGC CC |

Table S4: List of plasmids used

| Plasmid | Source | Reference |
|---|---|---|
| pCX-OKS-2A | Addgene plasmid # 19771 | [1] |
| FUGW | Addgene plasmid # 14883 | [2] |
| pBABE-cMycER | Dr. G. Evan | [3] |
| pLOVE-Klf4 | Addgene plasmid # 15950 | [4] |
| Mouse Nanog promoter luciferase reporter | Dr. T. Tada | [5] |
| psPAX2 | Addgene, cat# 12259 | Kind gift from Dr. Trono |
| pHCMV-EcoEnv | Addgene, cat# 15802 | [6] |

Table S5: List of qRT-PCR primers used in this study for mRNA expression analysis

| Name | Sequence |
|---|---|
| Ccna2_RT_F01 | ACA GAG TGT GAA GAT GCC CTG |
| Ccna2_RT_R01 | AAC GTT CAC TGG CTT GTC TTC |
| Cdh1_RT_F01 | GGC TGG ACC GAG AGA GTT AC |

| | |
|---|---|
| Cdh1_RT_R01 | CCG GGC ATT GAC CTC ATT CT |
| Cdh2_RT_F01 | GGG ACA TCA TCA CTG TGG CA |
| Cdh2_RT_R01 | TCC GTA GAA AGT CAT GGC AGT |
| Dppa3_RT_F01 | AGA CTT GTT CGG ATT GAG CAG A |
| Dppa3_RT_R01 | AAT GGC TCA CTG TCC CGT TC |
| GAPDH_RT_F01 | ATC AAC GGG AAG CCC ATC AC |
| GAPDH_RT_R01 | CCT TTT GGC TCC ACC CTT CA |
| Klf4_RT_F01 | ACA TTA ATG AGG CAG CCA CCT G |
| Klf4_RT_R01 | AGA GAG TTC CTC ACG CCA AC |
| Nanog_RT_F01 | TGA GCT ATA AGC AGG TTA AGA CC |
| Nanog_RT_R01 | CTG GGA TAC TCC ACT GGT GC |
| Oct_RT_Endo_F01 | GTG AGC CGT CTT TCC ACC AG |
| Oct_RT_Endo_R01 | ATA CCT CTG AGC CTG GTC CGA TTC CA |
| Oct_RT_Total_F01 | GTG GAG GAA GCC GAC AAC AAT GA |
| Oct_RT_Total_R01 | CAA GCT GAT TGG CGA TGT GAG |
| Snai1_RT_F01 | AAC TAT AGC GAG CTG CAG GA |
| Snai1_RT_R01 | GTA CCA GGA GAG AGT CCC AGA |
| Snai2_RT_F01 | AGA AGC CCA ACT ACA GCG AA |
| Snai2_RT_R01 | ATA GGG CTG TAT GCT CCC GA |
| Sox2_RT_F01 | TTT GTC CGA GAC CGA GAA GC |
| Sox2_RT_R01 | CTC CGG GAA GCG TGT ACT TA |
| Thy1_RT_F01 | GGA GTC CAG AAT CCA AGT CGG |
| Thy1_RT_R01 | TAT TCT CAT GGC GGC AGT CC |

Table S6: List of ChIP primers

| | |
|---|---|
| Ccna2_ChIP_AF | TCT CGA CAG CTA CGA CCA GAA CAC A |
| Ccna2_ChIP_AR | ACA TGT GGG ACG TCC ATC TGA GGT |
| Ccna2_ChIP_BF | GTC ATT CAG ATC CAT ACG CTC CTG CC |
| Ccna2_ChIP_BR | TGC TGC TCA GTG GAT CTG TAG CCT |
| Ccna2_ChIP_CF | AGC CTG CCT TCA CCA TTC ATG TGG |
| Ccna2_ChIP_CR | TGT CAG TGG TTT TCT TGC TCC GGG |

Table S7: List of Antibodies used

| Antibody | Catalogue No | Company | Assay | Dilution |
|---|---|---|---|---|
| Oct-3/4 | SC-8628 | Santa Cruz | WB<br>IF<br>ChIP | 1:600<br>1:100<br>5 µg |
| Klf4 | AF3158 | R & D | ChIP | 5 µg |
| Lamin B1 | ab16048 | Abcam | WB | 1:10,000 |
| Gapdh | ab8245 | Abcam | WB | 1:10,000 |

**Supplementary References:**

[1] K. Okita, M. Nakagawa, H. Hyenjong, T. Ichisaka, S. Yamanaka, Generation of mouse induced pluripotent stem cells without viral vectors, Science, 322 (2008) 949-953.

[2] C. Lois, E.J. Hong, S. Pease, E.J. Brown, D. Baltimore, Germline transmission and tissue-specific expression of transgenes delivered by lentiviral vectors, Science, 295 (2002) 868-872.

[3] T.D. Littlewood, D.C. Hancock, P.S. Danielian, M.G. Parker, G.I. Evan, A modified oestrogen receptor ligand-binding domain as an improved switch for the regulation of heterologous proteins, Nucleic acids research, 23 (1995) 1686-1690.

[4] R. Blelloch, M. Venere, J. Yen, M. Ramalho-Santos, Generation of induced pluripotent stem cells in the absence of drug selection, Cell stem cell, 1 (2007) 245-247.

[5] T. Kuroda, M. Tada, H. Kubota, H. Kimura, S.Y. Hatano, H. Suemori, N. Nakatsuji, T. Tada, Octamer and Sox elements are required for transcriptional cis regulation of Nanog gene expression, Mol Cell Biol, 25 (2005) 2475-2485.

[6] M. Sena-Esteves, J.C. Tebbets, S. Steffens, T. Crombleholme, A.W. Flake, Optimized large-scale production of high titer lentivirus vector pseudotypes, Journal of virological methods, 122 (2004) 131-139.
